# Cis-regulatory elements explain most of the mRNA stability variation across genes in yeast

**DOI:** 10.1101/085522

**Authors:** Jun Cheng, Kerstin C. Maier, Žiga Avsec, Petra Rus, Julien Gagneur

## Abstract

The stability of mRNA is one of the major determinants of gene expression. Although a wealth of sequence elements regulating mRNA stability has been described, their quantitative contributions to half-life are unknown. Here, we built a quantitative model for *Saccharomyces cerevisiae* based on functional mRNA sequence features that explains 60% of the half-life variation between genes and predicts half-life at a median relative error of 30%. The model revealed a new destabilizing 3’UTR motif, ATATTC, which we functionally validated. Codon usage proves to be the major determinant of mRNA stability. Nonetheless, single-nucleotide variations have the largest effect when occurring on 3’UTR motifs or upstream AUGs. Analyzing mRNA half-life data of 34 knockout strains showed that the effect of codon usage not only requires functional decapping and deadenylation, but also the 5’-to-3’ exonuclease Xrn1, the non-sense mediated decay genes, but not no-go decay. Altogether, this study quantitatively delineates the contributions of mRNA sequence features on stability in yeast, reveals their functional dependencies on degradation pathways, and allows accurate prediction of half-life from mRNA sequence.

## INTRODUCTION

The stability of messenger RNAs is an important aspect of gene regulation. It influences the overall cellular mRNA concentration, as mRNA steady-state levels are the ratio of synthesis and degradation rate. Moreover, low stability confers high turnover to mRNA and therefore the capacity to rapidly reach a new steady-state level in response to a transcriptional trigger (Shalem *et al*, 2008). Hence, stress genes, which must rapidly respond to environmental signals, show low stability (Zeisel *et al*, 2011; Miller *et al*, 2011; Rabani *et al*, 2014; Marguerat *et al*, 2014). In contrast, high stability provides robustness to variations in transcription. Accordingly, a wide range of mRNA-half-lives is observed in eukaryotes, with typical variations in a given genome spanning one to two orders of magnitude (Schwanhäusser *et al*, 2011; Schwalb *et al*, 2016; Eser *et al*, 2016). Also, significant variability in mRNA half-life among human individuals could be demonstrated for about a quarter of genes in lymphoblastoid cells and estimated to account for more than a third of the gene expression variability (Duan *et al*, 2013).

How mRNA stability is encoded in a gene sequence has long been a subject of study. Cis-regulatory elements (CREs) affecting mRNA stability are mainly encoded in the mRNA itself. They include but are not limited to secondary structure (Rabani *et al*, 2008; Geisberg *et al*, 2014), sequence motifs present in the 3’UTR including binding sites of RNA-binding proteins (Olivas & Parker, 2000; Shalgi *et al*, 2005; Duttagupta *et al*, 2005; Hogan *et al*, 2008; Hasan *et al*, 2014), and, in higher eukaryotes, microRNAs (Lee *et al*, 1993). Moreover, translation-related features are frequently associated with mRNA stability. For instance, inserting strong secondary structure elements in the 5’UTR or modifying the translation start codon context strongly destabilizes the long-lived *PGK1* mRNA in *S. cerevisiae* (Muhlrad *et al*, 1995; LaGrandeur & Parker, 1999). Codon usage, which affects translation elongation rate, also regulates mRNA stability (Hoekema *et al*, 1987; Presnyak *et al*, 2015; Mishima & Tomari, 2016; Bazzini *et al*, 2016), Further correlations between codon usage and mRNA stability have been reported in *E. coli* and *S. pombe* (Boël *et al*, 2016; Harigaya & Parker, 2016).

Since the RNA degradation machineries are well conserved among eukaryotes, the pathways have been extensively studied using *S. cerevisiae* as a model organism (Garneau *et al*, 2007; Parker, 2012). The general mRNA degradation pathway starts with the removal of the poly(A) tail by the Pan2/Pan3 (Brown *et al*, 1996) and Ccr4/Not complexes (Tucker *et al*, 2001). Subsequently, mRNA is subjected to decapping carried out by Dcp2 and promoted by several factors including Dhh1 and Pat1 (Pilkington & Parker, 2008; She *et al*, 2008). The decapped and deadenylated mRNA can be rapidly degraded in the 3’ to 5’ direction by the exosome (Anderson & Parker, 1998) or in the 5’ to 3’ direction by Xrn1 (Hsu & Stevens, 1993). Further mRNA degradation pathways are triggered when aberrant translational status is detected, including Nonsense-mediated decay (NMD), No-go decay (NGD) and Non-stop decay (NSD) (Garneau *et al*, 2007; Parker, 2012).

Despite all this knowledge, prediction of mRNA half-life from a gene sequence is still not established. Moreover, most of the mechanistic studies so far were only performed on individual genes or reporter genes. It is therefore unclear how the measured effects generalize genome-wide. A recent study showed that translation-related features can be predictive for mRNA stability (Neymotin *et al*, 2016). Although this analysis supported the general correlation between translation and stability (Edri & Tuller, 2014; Lackner *et al*, 2007), the model was not based purely on sequence-derived features. It also contained measured transcript properties such as ribosome density and normalized translation efficiencies. Hence, the question of how half-life is genetically encoded in mRNA sequence remains to be addressed.

Additionally, the dependencies of sequence features to distinct mRNA degradation pathways have not been systematically studied. One example of this is codon-mediated stability control. Although a causal link from codon usage to mRNA half-life has been shown for a wide range of organisms (Hoekema *et al*, 1987; Presnyak *et al*, 2015; Mishima & Tomari, 2016; Bazzini *et al*, 2016), the underlying mechanism remains poorly understood. In *S. cerevisiae*, reporter gene experiments showed that codon-mediated stability control depends on the RNA helicase Dhh1 (Radhakrishnan *et al*, 2016). However, it is unclear whether this generalizes to all mRNAs genome-wide. Also, the role of other closely related degradation pathways has not been systematically assessed with genome-wide half-life data.

Here, we mathematically modeled mRNA half-life as a function of its sequence. Applied to *S. cerevisiae*, our model can explain most of the between-gene half-life variance from sequence alone. Using a semi-mechanistic model, we could interpret individual sequence features in the 5’UTR, coding region, and 3’UTR. Quantification of the respective contributions revealed that codon usage is the major contributor to mRNA stability. Applying the modeling approach to *S. pombe* supports the generality of these findings. Moreover, we systematically assessed the dependencies of these sequence elements on mRNA degradation pathways using half-life data for 34 knockout strains. This revealed in particular novel pathways through which codon usage affects half-life.

## RESULTS

To study cis-regulatory determinants of mRNA stability in *S. cerevisiae*, we chose the dataset by Sun and colleagues (Sun *et al*, 2013) which provides genome-wide half-life measurements for 4,388 expressed genes of a wild-type lab strain and 34 strains knocked out for RNA degradation pathway genes (Figure 1, Supplementary Table 1). When applicable, we also investigated half-life measurements of *S. pombe* for 3,614 expressed mRNAs in a wild-type lab strain from Eser and colleagues (Eser *et al*, 2016). We considered sequence features within 5 overlapping regions: the 5’UTR, the start codon context, the coding sequence, the stop codon context and the 3’UTR. We assessed their effects in the wild type and in the 34 knockout strains (Figure 1). Finally, we fitted a joint model to assess the contribution of individual sequence features and their single-nucleotide effects (Figure 1).

**Figure 1:**
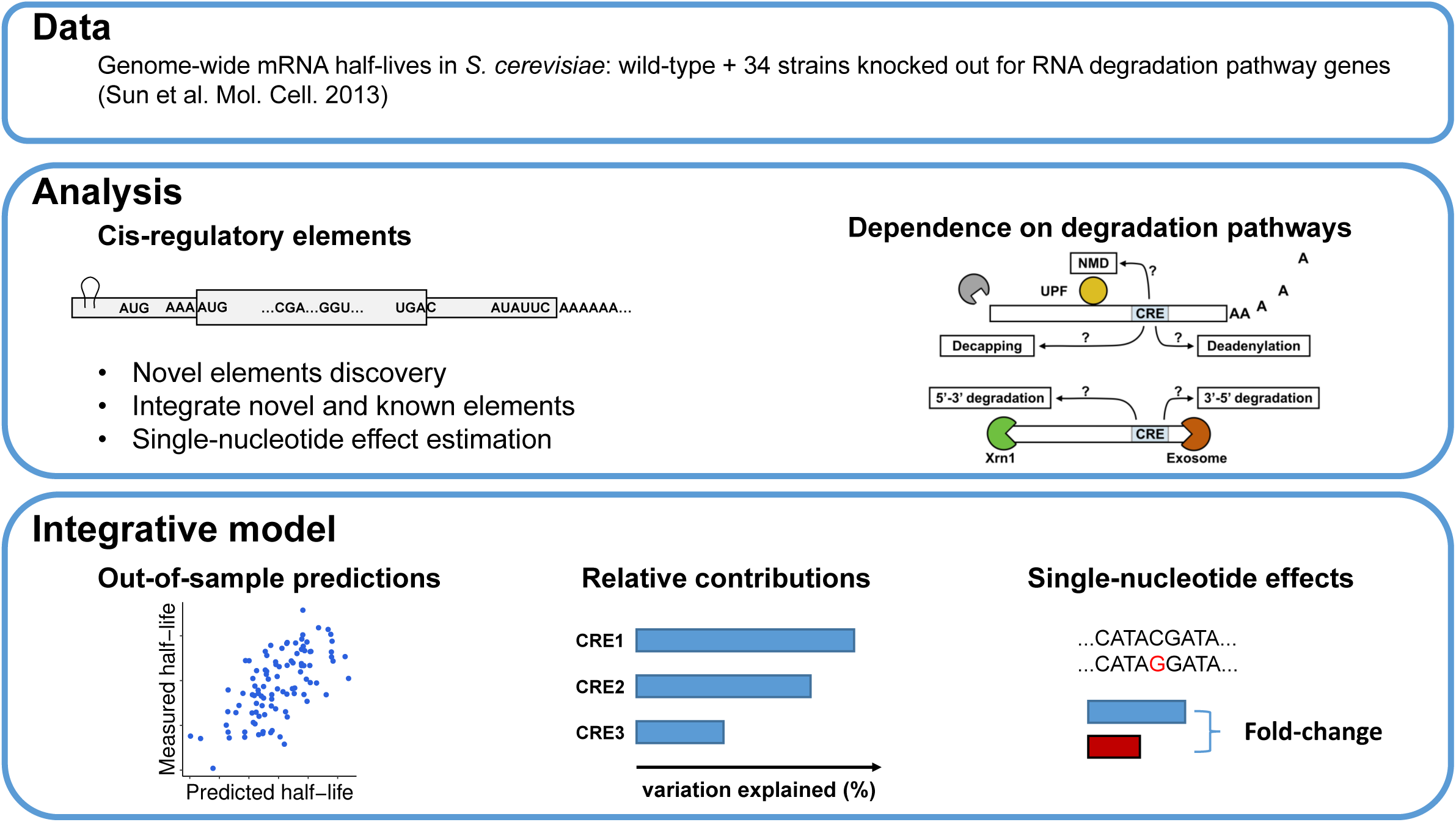
Study overview. The goal of this study is to discover and integrate cis-regulatory mRNA elements affecting mRNA stability and assess their dependence on mRNA degradation pathways. **Data)** we obtained *S. cerevisiae* genome-wide half-life data from wild-type (WT) as well as from 34 knockout strains from Sun et al. (2013). Each of the knockout strains has one gene closely related to mRNA degradation pathways knocked out. **Analysis)** we systematically searched for novel sequence features associating with half-life from 5*’*UTR, start codon context, CDS, stop codon context, and 3*’*UTR. Effects of previously reported cis-regulatory elements were also assessed. Moreover, we assessed the dependencies of different sequence features on degradation pathways by analyzing their effects in the knockout strains. **Integrative model)** we build a statistical model to predict genome-wide half-life solely from mRNA sequence. This allowed the quantification of the relative contributions of the sequence features to the overall variation across genes and assessing the sensitivity of mRNA stability with respect to single-nucleotide variants.

The correlations between sequence lengths, GC contents and folding energies (Materials and Methods) with half-life and corresponding P-values are summarized in Supplementary Table 2 and Supplementary Figures S1-3. In general, sequence lengths correlated negatively with half-life and folding energies correlated positively with half-life in both yeast species, whereas correlations of GC content varied with species and gene regions.

In the following subsections, we describe first the findings for each of the 5 gene regions and then a model that integrates all these sequence features.

### Upstream AUGs destabilize mRNAs by triggering nonsense-mediated decay

Occurrence of an upstream AUG (uAUG) associated significantly with shorter half-life (median fold-change = 1.37, *P* < 2 × 10^−16^). This effect was strengthened for genes with two or more AUGs (Figure 2A,B). Among the 34 knock-out strains, the association between uAUG and shorter half-life was almost lost only for mutants of the two essential components of the nonsense-mediated mRNA decay (NMD) *UPF2* and *UPF3* (Leeds *et al*, 1992; Cui *et al*, 1995), and for the general 5’-3’ exonuclease *Xrn1* (Figure 2A). The dependence on NMD suggested that the association might be due to the occurrence of a premature stop codon. Consistent with this hypothesis, the association of uAUG with decreased half-lives was only found for genes with a premature stop codon cognate with the uAUG (Figure 2C). This held not only for cognate premature stop codons within the 5’UTR, leading to a potential upstream ORF, but also for cognate premature stop codons within the ORF, which occurred almost always for uAUG out-of-frame with the main ORF (Figure 2C). This finding likely holds for many other eukaryotes as we found the same trends in *S. pombe* (Figure 2D). These observations are consistent with a single-gene study demonstrating that translation of upstream ORFs can lead to RNA degradation by nonsense-mediated decay (Gaba *et al*, 2005). Altogether, these results show that uAUGs are mRNA destabilizing elements as they almost surely match with cognate premature stop codons, which, whether in frame or not with the gene, and within the UTR or in the coding region, trigger NMD.

**Figure 2:**
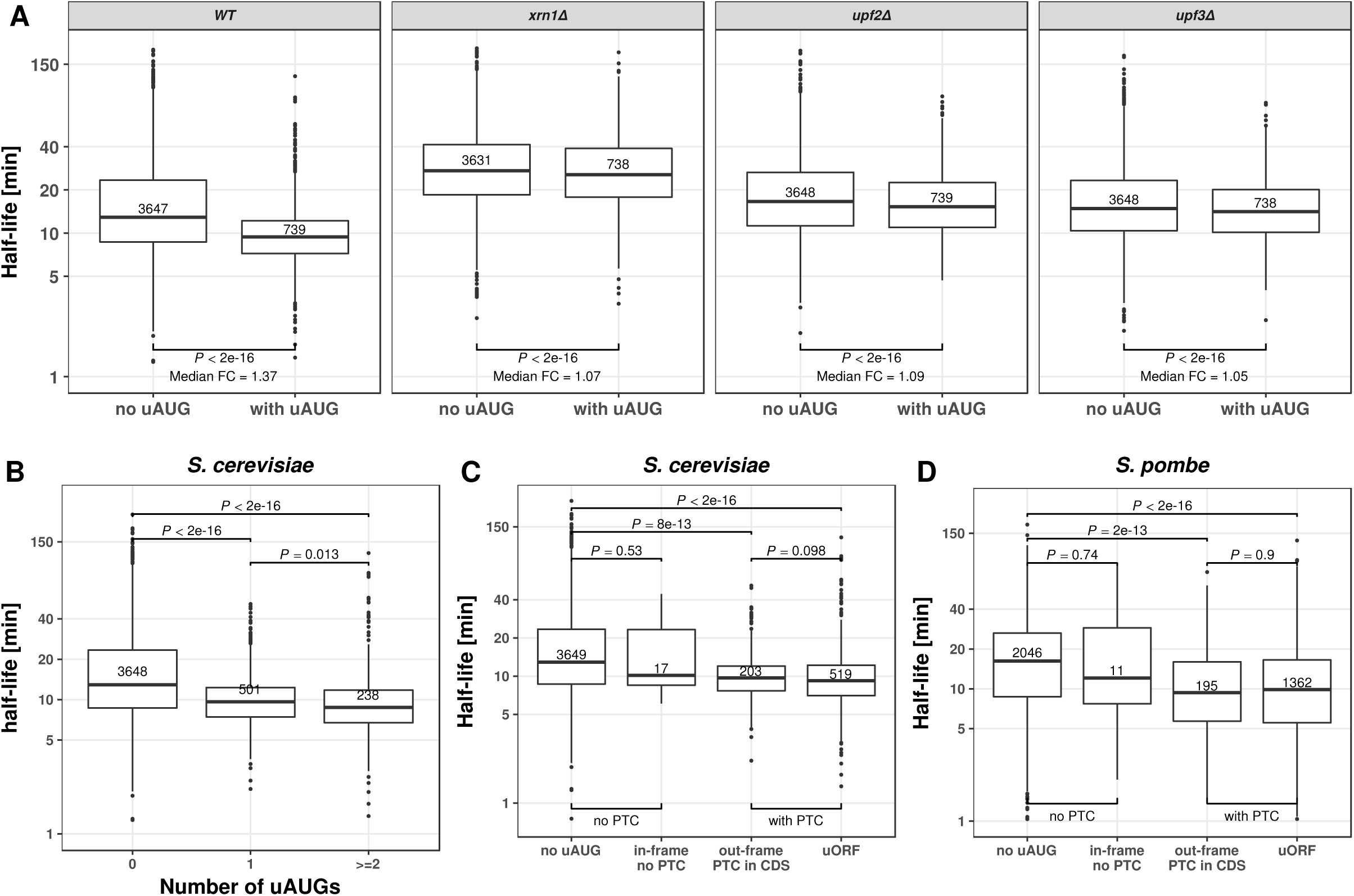
Upstream AUG codon (uAUG) destabilize mRNA. (**A**) Distribution of mRNA half-life for mRNAs without uAUG eft) and with at least one uAUG (right) in, from left to right: wild type, *XRN1*, *UPF2* and *UPF3* knockout *S. cerevisiae* strains. Median fold-change (Median FC) calculated by dividing the median of the group without uAUG with the group with uAUG. (**B**) Distribution of mRNA half-lives for mRNAs with zero (left), one (middle), or more (right) uAUGs in *S. cerevisiae*. (**C**) Distribution of mRNA half-lives for *S. cerevisiae* mRNAs with, from left to right: no uAUG, with one in-frame uAUG but no cognate premature termination codon, with one out-of-frame uAUG and one cognate premature termination codon in the CDS, and with one uAUG and one cognate stop codon in the 5*’*UTR (uORF). (**D**) Same as in (C) for *S. pombe* mRNAs. All p-values were calculated with Wilcoxon rank-sum test. Numbers in the boxes indicate number of members in the corresponding group. Boxes represent quartiles, whiskers extend to the highest or lowest value within 1.5 times the interquartile range and horizontal bars in the boxes represent medians. Data points falling further than 1.5-fold the interquartile distance are considered outliers and are shown as dots.

### Translation initiation sequence features associate with mRNA stability

Several sequence features in the 5’UTR including the start codon context associated with mRNA half-life (Expanded view, Supplementary Figure S4-5). This indicates that 5’UTR elements may affect mRNA stability by altering translation initiation. However, none of these sequence features remained significant in the final joint model. Our analysis is therefore not conclusive on this point. A detailed analysis is provided in the Expanded View for the interested readers.

### Codon usage regulates mRNA stability through common mRNA decay pathways

Codon usage marginally explained 55% of the between-gene half-life variation in *S. cerevisiae* (Figure 3A). Species-specific tRNA adaptation index (sTAI) (Sabi & Tuller, 2014) significantly correlated with half-life in both *S. cerevisiae* (Supplementary Figure S4E, *ρ* = 0.55, *P* < 2.2x10^−16^) and *S. pombe* (Supplementary Figure S4F, *ρ* = 0.41, *P* < 2. 2x10^−16^), confirming previously observed association between codon optimality and mRNA stability (Presnyak *et al*, 2015; Harigaya & Parker, 2016). Next, using the out-of-folds explained variance as a summary statistics, we assessed its variation across different gene knockouts (Materials and Methods). The effect of codon usage exclusively depended on the genes from the common deadenylation- and decapping-dependent 5’ to 3’ mRNA decay pathway and the NMD pathway (all FDR < 0.1, Figure 3B). In particular, all assayed genes of the Ccr4-Not complex, including *CCR4*, *NOT3*, *CAF40* and *POP2*, were required for wild-type level effects of codon usage on mRNA decay. Among them, *CCR4* has the largest effect. This confirmed a recent study in zebrafish showing that accelerated decay of non-optimal codon genes requires deadenylation activities of Ccr4-Not (Mishima & Tomari, 2016). In contrast to genes of the Ccr4-Not complex, *PAN2/3* genes which encode also deadenylation enzymes, were not found to be essential for the coupling between codon usage and mRNA decay (Figure 3B).

**Figure 3:**
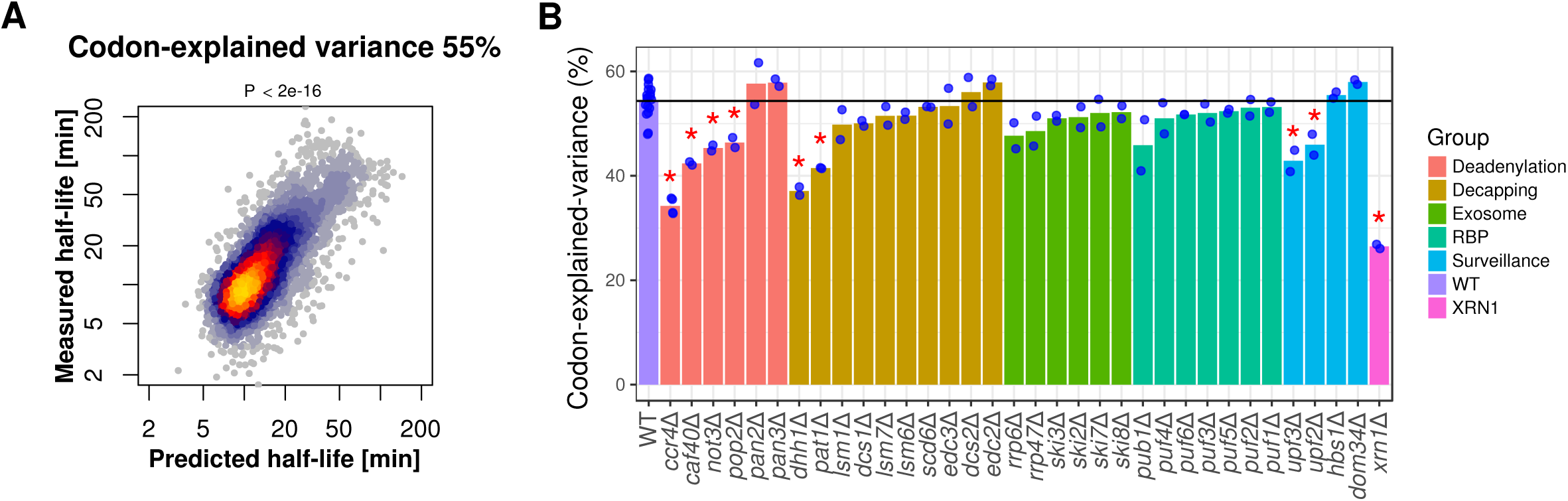
Codon usage regulates mRNA stability through common mRNA decay pathways. (**A**) Predicted mRNA half-life using only codons as features (linear regression) versus measured mRNA half-life. (**B**) mRNA half-life explained variance (y-axis, Materials and Methods) in wild-type (WT) and across all 34 knockout strains (grouped according to their functions). Each blue dot represents one replicate, bar heights indicate means across replicates. Bars with a red star are significantly different from the wild type level (FDR <0.1, Wilcoxon rank-sum test, followed by Benjamini-Hochberg correction).

Furthermore, our results not only confirm the dependence on Dhh1 (Radhakrishnan *et al*, 2016), but also on its interacting partner Pat1. Our findings of Pat1 and Ccr4 contradict the negative results for these genes reported by Radhakrishnan *et al.* (Radhakrishnan *et al*, 2016). The difference might come from the fact that our analysis is genome-wide, whereas Radhakrishnan and colleagues used a reporter assay.

Our systematic analysis revealed two additional novel dependencies: First, on the common 5’ to 3’ exonuclease Xrn1, and second, on *UPF2 and UPF3* genes, which are essential players of NMD (all FDR < 0.1, Figure 3B). Previous studies have shown that NMD is more than just a RNA surveillance pathway, but rather one of the general mRNA decay mechanisms that target a wide range of mRNAs, including aberrant and normal ones (He *et al*, 2003; Hug *et al*, 2015). Notably, we did not observe any change of effect upon knockout of *DOM34* and *HBS1* (Supplementary Figure 6), which are essential genes for the No-Go decay pathway. This implies that the effect of codon usage is unlikely due to stalled ribosomes at non-optimal codons.

Altogether, our analysis strongly indicates that, the so-called “codon-mediated decay” is not an mRNA decay pathway itself, but a regulatory mechanism of the common mRNA decay pathways.

### Stop codon context associates with mRNA stability

The first nucleotide 3’ of the stop codon significantly associated with mRNA stability. This association was observed for each of the three possible stop codons, and for each codon a cytosine significantly associated with lower half-life (Supplementary Figure S4, also for P-values and fold changes). However, this feature was not significant in the joint model, and analysis of the knockout strains did not reveal clear pathway dependencies for it (Supplementary Figure S6). Detailed description is provided in the Expanded View for interested readers.

### Sequence motifs in 3’UTR

De novo motif search identified four motifs in the 3’UTR to be significantly associated with mRNA stability (Figure 4A, Materials and Methods). These include three described motifs: the Puf3 binding motif TGTAAATA (FDR = 3.2x10^−5^, median fold-change 1.29) (Gerber *et al*, 2004; Gupta *et al*, 2014), the Whi3 binding motif TGCAT (FDR = 7x10^−4^, median fold-change 1.24) (Colomina *et al*, 2008; Cai & Futcher, 2013), and a poly(U) motif TTTTTTA (FDR = 0.09, median fold-change 1.20), which can be bound by Pub1 (Duttagupta *et al*, 2005), or is part of the long poly(U) stretch that forms a looping structure with poly(A) tail (Geisberg *et al*, 2014). Moreover, an uncharacterized motif, ATATTC, was associated with lower mRNA half-life (FDR = 2x10^−5^, median fold-change 1.24). Genes harboring the ATATTC motif are significantly enriched for genes involved in oxidative phosphorylation (Bonferroni corrected *P* < 0.01, 4.4 fold enrichment, Gene Ontology analysis, Supplementary Methods and Supplementary Table 3). The motif ATATC preferentially localizes in the vicinity of the poly(A) site (Figure 4B), and functionally depends on Ccr4 (FDR < 0.1, Supplementary Figure 6), suggesting a potential interaction with deadenylation factors. This 3’UTR motif had been computationally identified by conservation analysis (Kellis *et al*, 2003), by regression of steady-state expression levels (Foat *et al*, 2005), and by enrichment analysis within genes expression clusters (Elemento *et al*, 2007). The motif was suggested to be named as PRSE (Positive Response to Starvation Element), because of its enrichment among genes up-regulated upon starvation (Foat *et al*, 2005). However, it was not experimentally validated for controlling of mRNA stability.

**Figure 4:**
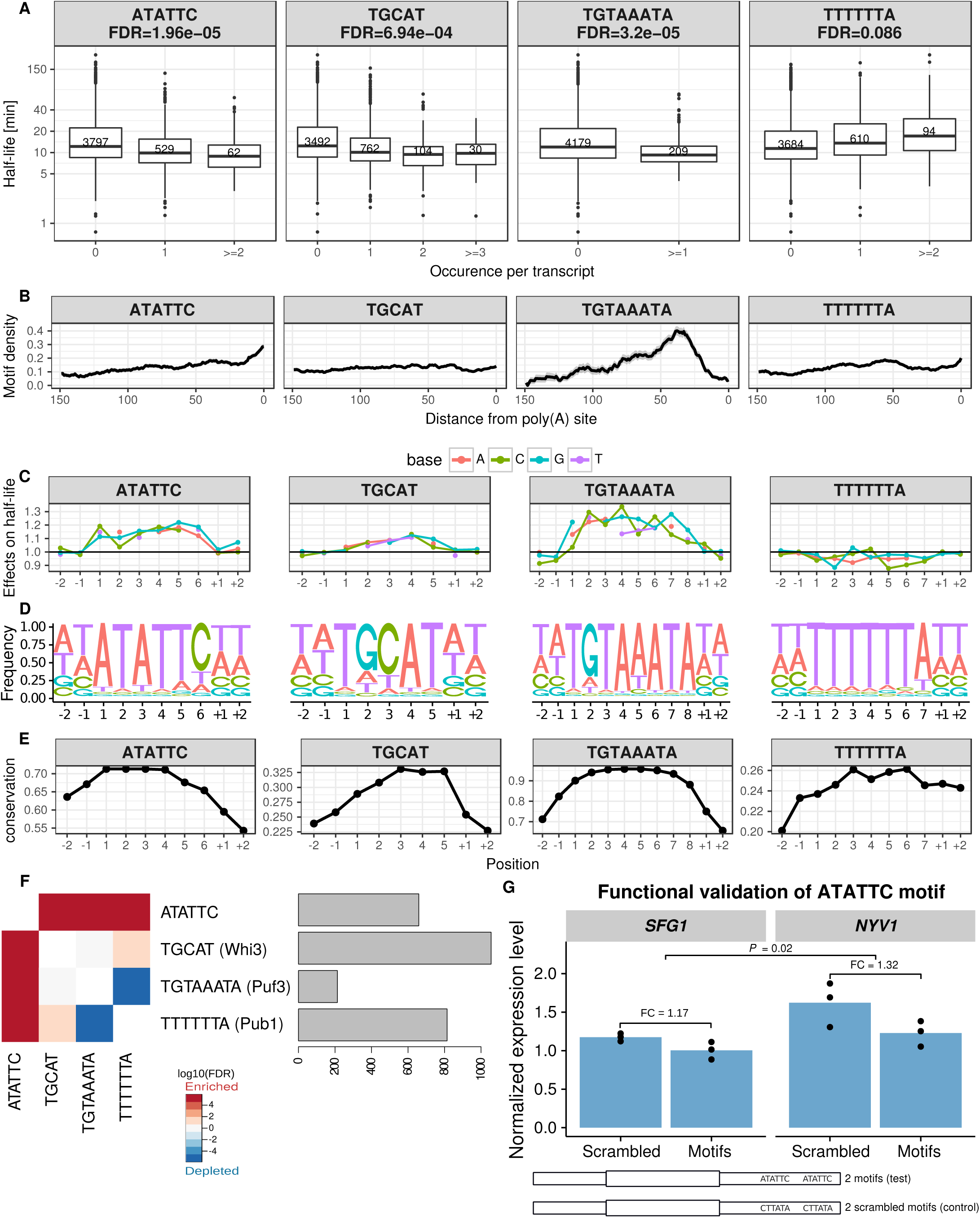
3’UTR half-life determinant motifs in *S. cerevisiae*. (**A**) Distribution of half-lives for mRNAs grouped by the number of occurrence(s) of the motif ATATTC, TGCAT (Whi3), TGTAAATA (Puf3) and TTTTTTA (Pub1) respectively in their 3’UTR sequence. Numbers in the boxes represent the number of members in each box. FDR were reported from the linear mixed effect model (Materials and Methods). (**B**) Fraction of transcripts containing the motif (y-axis) within a 20-bp window centered at a position (x-axis) with respect to poly(**A**) site for different motifs (facet titles). Positional bias was not observed when ligning 3’UTR motifs with respect to the stop codon. (**C**) Prediction of the relative effect on half-life (y-axis) for single-nucleotide substitution in the motif with respect to the consensus motif (y=1, horizontal line). The motifs were extended 2 bases at each anking site (positions +1, +2, -1, -2). (D) Nucleotide frequency within motif instances, when allowing for one mismatch compared to the consensus motif. (**E**) Mean conservation score (phastCons, Materials and Methods) of each base in the consensus motif with 2 anking nucleotides (y-axis). (**F**) Co-occurrence significance (FDR, Fisher test p-value corrected with Benjamini-Hochberg) between different motifs (left). Number of occurrences among the 4,388 mRNAs (right). (**G**) Steady-state expression level of *SFG1* and *NYV1* (normalized by *ACT1* and *TUB2* expression, Supplementary Methods). P-value calculated by joint comparing normalized expression level of constructs that with 2 scrambled motifs embedded versus that with 2 functional ATATTC motifs embedded (Supplementary Methods).

We validated the 3’UTR motif ATATTC with a reporter assay on two different genes *SFG1* and *NYV1*. Given the predicted small effect of a single motif, we generated constructs with two instances of the motif and compared them to constructs harboring two scrambled motifs at the same locations (Figure 4G, Material and Methods). Both reporter genes showed decreased expression levels compared to scrambled controls (P = 0.02, mean fold-change 1.25, Supplementary Methods). Since the 3’UTR motif ATATTC motif is not significantly associated with mRNA synthesis rate (*P* = 0.38, Wilcoxon rank-sum test, half-life of genes without motif versus genes with motif), we conclude that this decreased expression is due to decreased stability.

Consistent with the role of Puf3 in recruiting deadenylation factors, Puf3 binding motif localized preferentially close to the poly(A) site (Figure 4B). The effect of the Puf3 motifs was significantly lower in the knockout of *PUF3* (FDR < 0.1, Supplementary Figure 6). We also found a significant dependence on the deadenylation (*CCR4*, *POP2*) and decapping (*DHH1*, *PAT1*) pathways (all FDR < 0.1, Supplementary Figure 6), consistent with previous single gene experiment showing that Puf3 binding promotes both deadenylation and decapping (Olivas & Parker, 2000; Goldstrohm *et al*, 2007). Strikingly, Puf3 binding motif switched to a stabilization motif in the absence of Puf3 and Ccr4 (all FDR < 0.1, Supplementary Figure 6), suggesting that deadenylation of Puf3 motif containing mRNAs is not only facilitated by Puf3 binding, but also depends on it.

Whi3 plays an important role in cell cycle control (Garí *et al*, 2001). Binding of Whi3 leads to destabilization of the *CLN3* mRNA (Cai & Futcher, 2013). A subset of yeast genes are up-regulated in the Whi3 knockout strain (Cai & Futcher, 2013). However, it was so far unclear whether Whi3 generally destabilizes mRNAs upon its binding. Our analysis showed that mRNAs containing the Whi3 binding motif (TGCAT) have significantly shorter half-life (FDR = 6.9x10^−04^, median fold-change 1.24). Surprisingly, this binding motif is extremely widespread, with 896 out of 4,388 (20%) genes that we examined containing the motif on the 3’UTR region, which enriched for genes involved in several processes (Supplementary Table 3). Functionality of Whi3 binding motif was found to be dependent on Ccr4 (FDR < 0.1, Supplementary Figure 6).

The mRNAs harboring the TTTTTTA motif tended to be more stable and enriched for translation (*P* = 1.34x10^−03^, 2 fold enrichment, Supplementary Table 3, Figure 4A). No positional preferences were observed for this motif (Figure 4B). Effects of this motif depends on genes from Ccr4-Not complex and Xrn1 (Supplementary Figure 6).

Additional three lines of evidence further supported the functionality of our identified motifs. First, single nucleotide deviations from the motif’s consensus sequence associated with decreased effects on half-life (Figure 4C, linear regression allowing for one mismatch, Materials and Methods). Moreover, the flanking nucleotides did not show further associations indicating that the whole lengths of the motifs were recovered (Figure 4C). Second, when allowing for one mismatch, the motif still showed strong preferences (Figure 4D). Third, the motif instances were more conserved than their flanking bases from the 3’UTR (Figure 4E). Notably, the motif ATATTC, was found in 13% of the genes (591 out of 4,388) and significantly co-occurred with the other two destabilizing motifs found in 3’UTR: Puf3 motif (FDR = 0.01) and Whi3 motif (FDR = 3× 10^−3^) binding motifs (Fig 4F). Furthermore, all four motifs show significant effects in further RNA half-life datasets generated with distinct protocols (Supplementary Figure 7).

### 60% between-gene half-life variation can be explained by sequence features

We next asked how well one could predict mRNA half-life from these mRNA sequence features, and what their respective contributions were when considered jointly. To this end, we performed a multivariate linear regression of the logarithm of the half-life against the identified sequence features. The predictive power of the model on unseen data was assessed using 10-fold cross validation (Material and Methods, a complete list of model features and their p-values is provided in Supplementary Table 4). To prevent over-fitting, we performed motif discovery on each of the 10 training sets and observed the same set of motifs across all the folds. Altogether, 60% of *S*. *cerevisiae* half-life variance in the logarithmic scale can be explained by simple linear combinations of the above sequence features (Figure 5A, Supplementary Table 5). The median out-of-folds relative error across genes is 30%. A median relative error of 30% for half-life is remarkably low because it is in the order of magnitude of the expression variation that is typically physiologically tolerated, and it is also about the amount of variation observed between replicate experiments (Eser *et al*, 2016). To make sure that our findings are not biased to a specific dataset, we fitted the same model to a dataset using RATE-seq (Neymotin *et al*, 2014), a modified version of the protocol used by Sun and colleagues (Sun *et al*, 2013). On this data, the model was able to explain 50% of the variance (Supplementary Figure 8). Moreover, the same procedure applied to *S. pombe* explained 47% of the total half-life variance, suggesting the generality of this approach. Because the measures also entail measurement noise, these numbers are conservative underestimations of the total biological variance explained by our model.

**Figure 5:**
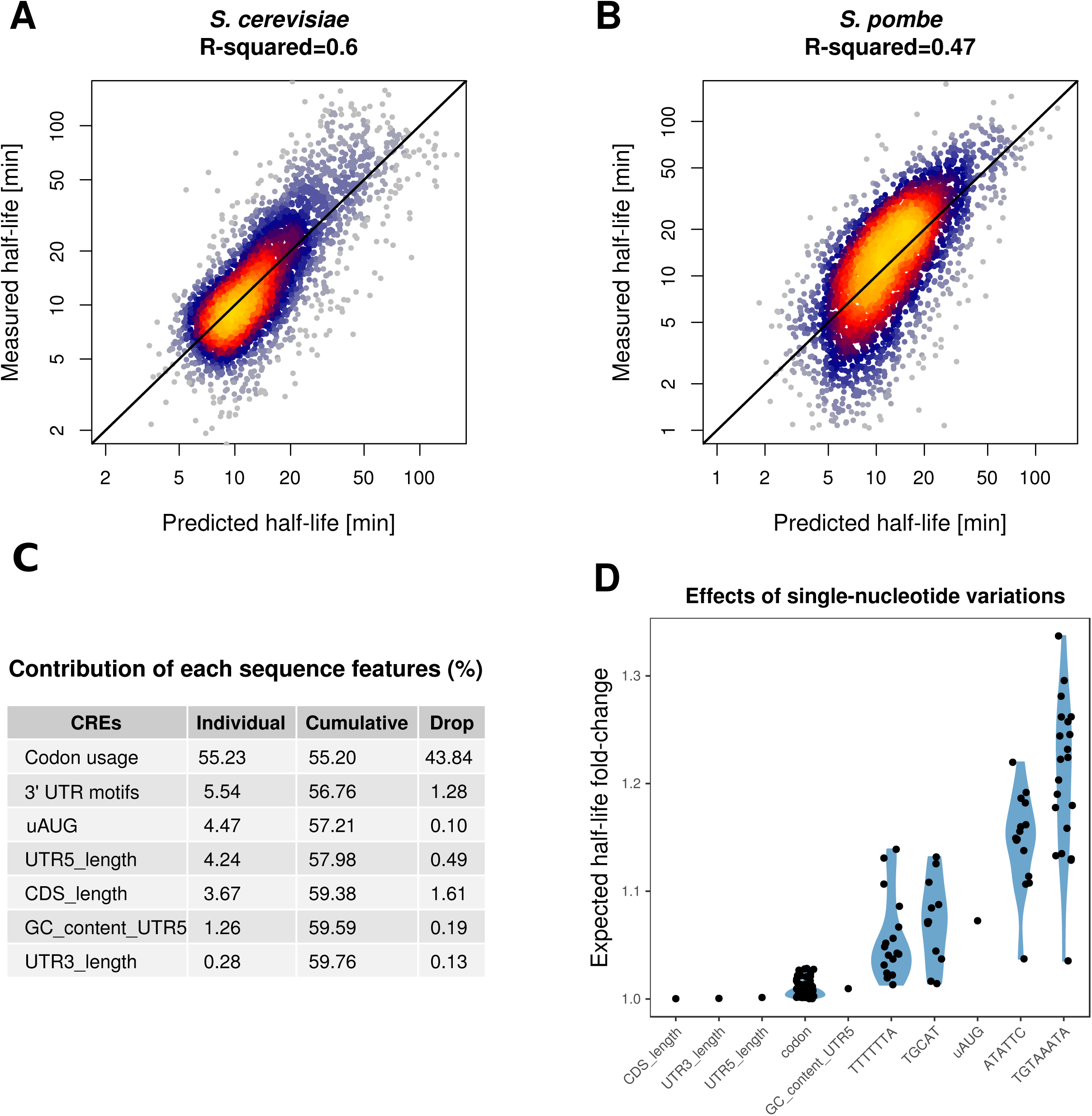
Genome-wide prediction of mRNA half-lives from sequence features and analysis of the contributions. **(A-B)** mRNA half-lives predicted (x-axis) versus measured (y-axis) for *S. cerevisiae* (A) and *S. pombe* (B) respectively. (**C**) Contribution of each sequence feature individually (Individual), cumulatively when sequentially added into a combined model (Cumulative) and explained variance drop when each single feature is removed from the full model separately (Drop). Values reported are the mean of 100 times of cross-validated evaluation (Materials and Methods). (**D**) Expected half-life fold-change of single-nucleotide variations on sequence features. For length and GC, dot represent median half-life fold change of one nucleotide shorter or one G/C to A/T transition respectively. For codon usage, each dot represents median half-life fold-change of one type of synonymous mutation, all kinds of synonymous mutations are considered. For uAUG, each dot represents median half-life fold-change of mutating out one uAUG. For motifs, each dot represents median half-life fold-change of one type of nucleotide transition at one position on the motif (Materials and Methods). Medians are calculated across all mRNAs.

**Figure 6:**
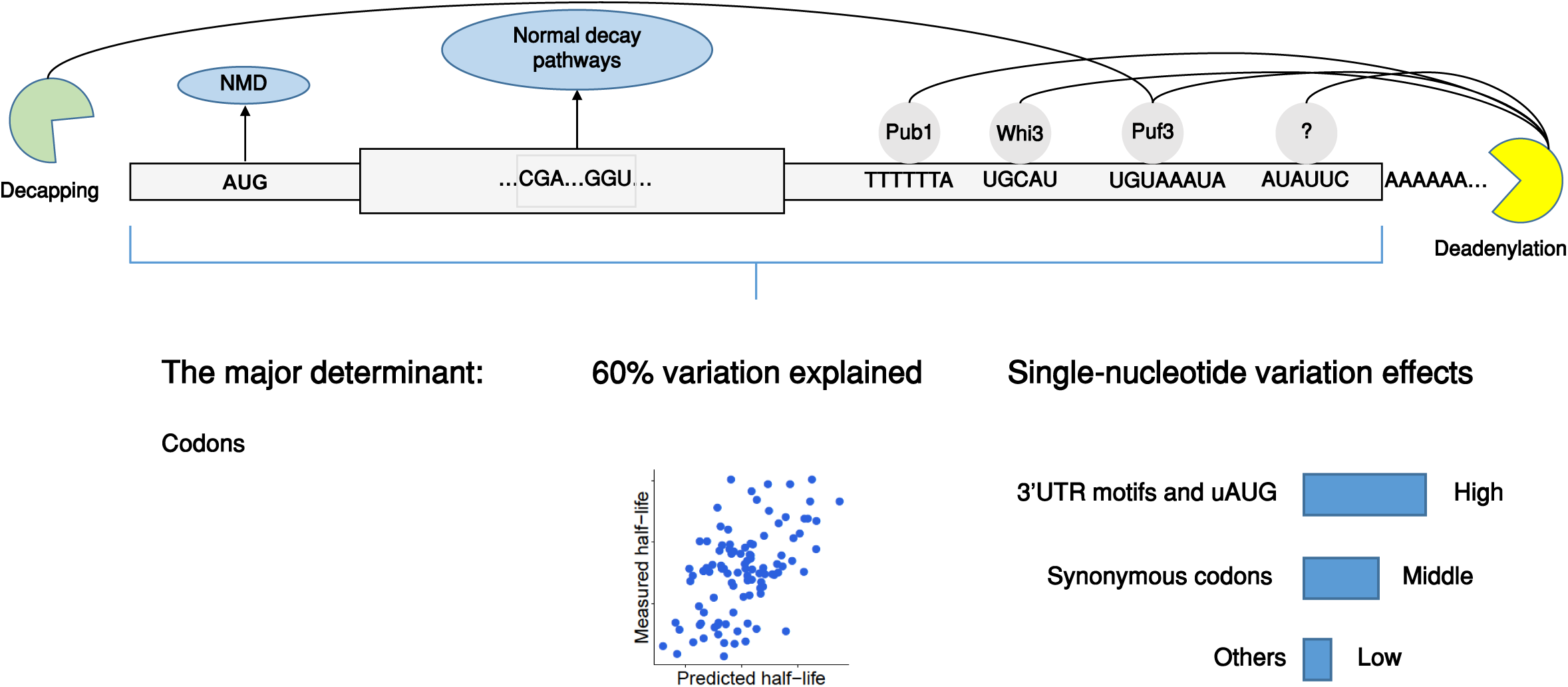
Overview of conclusions. Summary of conclusions from this study.

The uAUG, 5’UTR length, 5’UTR GC content, 61 coding codons, CDS length, all four 3’UTR motifs, and 3’UTR length remained significant in the joint model indicating that they contributed individually to half-life (complete list of p-values given in Supplementary Table 4). Most of them showed decreased effect in a joint model compared to marginal effects (Figure 5C), likely because they correlate with each other. In contrast, start codon context, stop codon context, 5’ folding energy, the 5’UTR motif AAACAAA (Supplementary Figure S5), and 3’UTR GC content dropped below the significance when considered in the joint model (Supplementary Table 4). This loss of statistical significance may be due to lack of statistical power. Another possibility is that the marginal association of these sequence features with half-life is a consequence of a correlation with other sequence features. Among all sequence features, codon usage as a group is the best predictor both in a univariate model (55.23%) and in the joint model (43.84 %) (Figure 5C). This shows that, quantitatively, codon usage is the major determinant of mRNA stability in yeast. This explains why only a small fraction of mRNA stability variation can be explained by RNA-binding proteins (Hasan *et al*, 2014). The variance analysis quantifies the contribution of each sequence feature to the variation across genes. Features that vary a lot between genes, such as UTR length and codon usage, favorably contribute to the variation. However, this does not reflect the effect on a given gene of elementary sequence variations in these features. For instance, a single-nucleotide variant can lead to the creation of an uAUG with a strong effect on half-life, but a single nucleotide variant in the coding sequence may have little impact on overall codon usage. We used the joint model to assess the sensitivity of each feature to single-nucleotide mutations as median fold-change across genes, simulating single-nucleotide deletions for the length features and single nucleotide substitutions for the remaining ones (Materials and Methods). Single-nucleotide variations typically altered half-life by less than 10%. The largest effects were observed in the 3’UTR motifs and uAUG (Figure 5D). Notably, although codon usage was the major contributor to the variance, synonymous variation on codons typically affected half-life by less than 2% (Figure 5D; Supplementary Figure 9). For those synonymous variations that changed half-life by more than 2%, most of them were variations that involved the most non-optimized codons CGA or ATA (Supplementary Figure 9, Presnyak et al. 2015).

Altogether, our results show that most of yeast mRNA half-life variation can be predicted from mRNA sequence alone, with codon usage being the major contributor. However, single-nucleotide variation at 3’UTR motifs or uAUG had the largest expected effect on mRNA stability.

## DISCUSSION

We systematically searched for mRNA sequence features associating with mRNA stability and estimated their effects at single-nucleotide resolution in a joint model. Up to GC content and length, all elements of the joint model are causal. One of them, the 3’UTR motif ATATTC has been validated in this study. Overall, the joint model showed that 60% of the variance could be predicted from mRNA sequence alone in *S. cerevisiae*. This analysis showed that translation-related features, in particular codon usage, contributed most to the explained variance. This findings strengthens further the importance of the coupling between translation and mRNA degradation (Roy & Jacobson, 2013; Huch & Nissan, 2014; Radhakrishnan & Green, 2016). Moreover, we assessed the dependencies of each sequence element on RNA degradation pathways. Remarkably, we identified that codon-mediated decay is a regulatory mechanism of the canonical decay pathways, including deadenylation- and decapping-dependent 5’ to 3’ decay and NMD (Figure 5E).

Predicting various steps of gene expression from sequence alone has long been a subject of study (Beer & Tavazoie, 2004; Vogel *et al*, 2010; Zur & Tuller, 2013; Wang *et al*, 2016). To this end, two distinct classes of models have been proposed: The biophysical models on the one hand and the machine learning models on the other hand (Zur & Tuller, 2016). Biophysical models provide detailed understanding of the processes. On the other hand, machine learning approaches can reach much higher predictive accuracy but are more difficult to interpret. Also, machine learning approaches can pick up signals with predictive power that are correlative but not causal. Here we adopted an intermediate, semi-mechanistic modeling approach. We used a simple linear model that is interpretable. Also, all elements are functional, up to two covariates: GC content and length.

Our approach was based on the analysis of endogenous sequence, which allowed the identification of a novel cis-regulatory element. An alternative approach to the modeling of endogenous sequence is to use large-scale synthetic libraries (Dvir *et al*, 2013; Shalem *et al*, 2015; Wissink *et al*, 2016). Although very powerful to dissect known cis-regulatory elements or to investigate small variations around select genes, the sequence space is so large that these large-scale perturbation screens cannot uncover all regulatory motifs. It would be interesting to combine both approaches and design large-scale validation experiments guided by insights coming from modeling of endogenous sequences as we developed here.

Recently, Neymotin and colleagues (Neymotin *et al*, 2016) showed that several translation-related transcript properties associated with half-life. This study derived a model explaining 50% of the total variance using many transcript properties including some not based on sequence (ribosome profiling, expression levels, etc.). Although non-sequence based predictors can facilitate prediction, they may do so because they are consequences rather than causes of half-life. For instance increased half-life causes higher expression level. Also, increased cytoplasmic half-life, provides a higher ratio of cytoplasmic over nuclear RNA, and thus more RNAs available to ribosomes. Hence both expression level and ribosome density may help making good predictions of half-life, but not necessarily because they causally increase half-life. In contrast, we aimed here to understand how mRNA half-life is encoded in mRNA sequence and derived a model that is based on functional elements. This avoided using transcript properties which could be consequences of mRNA stability. Hence, our present analysis confirms the quantitative importance of translation in determining mRNA stability that Neymotin and colleagues quantified, and anchors it into pure sequence elements.

Confounding associations of sequence elements with mRNA stability could arise because of selection on expression levels acting at multiple stage of gene expression. For instance, genes that are selected for high protein expression levels may be enriched for elements that enhance translation and for elements that enhance mRNA stability. Functional validations are therefore needed to disentangle causality from co-selection. The sequence elements of our joint model, up to GC content and length, are all functional. However, we reported further elements that associate marginally with half-life. One of the interesting sequence elements that we found associated with half-life but did not turn out significant in the joint model is the start codon context. Given its established effect on translation initiation (Dvir *et al*, 2013; Kozak, 1986), the general coupling between translation and mRNA degradation (Roy & Jacobson, 2013; Huch & Nissan, 2014; Radhakrishnan & Green, 2016), as well as several observations directly on mRNA stability for single genes (LaGrandeur & Parker, 1999; Schwartz & Parker, 1999), the start codon context may nonetheless functionally affect mRNA stability. Consistent with this hypothesis, large scale experiments that perturb 5’ sequence secondary structure and start codon context indeed showed a wide range of mRNA level changes in the direction that we would predict (Dvir *et al*, 2013).

We are not aware of previous studies that systematically assessed the effects of cis-regulatory elements in the context of knockout backgrounds, as we did here. This part of our analysis turned out to be very insightful. By assessing the dependencies of codon usage mediated mRNA stability control systematically and comprehensively, we generalized results from recent studies on the Ccr4-Not complex and Dhh1, but also identified important novel ones including NMD factors, Pat1 and Xrn1. With the growing availability of knockout or mutant background in model organisms and human cell lines, we anticipate this approach to become a fruitful methodology to unravel regulatory mechanisms.

## MATERIALS AND METHODS

### Data and Genomes

Wild-type and knockout genome-wide *S. cerevisiae* half-life data were obtained from Sun and colleagues (Sun *et al*, 2013), whereby all strains are histidine, leucine, methionine and uracil auxotrophs. A complete list of knockout strains used in this study is provided in Supplementary Table 1. *S. cerevisiae* gene boundaries were taken from the boundaries of the most abundant isoform quantified by Pelechano and colleagues (Pelechano *et al*, 2013). Reference genome fasta file and genome annotation were obtained from the Ensembl database (release 79). UTR regions were defined by subtracting out gene body (exon and introns from the Ensembl annotation) from the gene boundaries. Processed *S. cerevisiae* UTR annotation is provided at Supplementary Table 6.

Genome-wide half-life data of *S. pombe* as well as refined transcription unit annotation were obtained from Eser and colleagues (Eser *et al*, 2016). Reference genome version ASM294v2.26 was used to obtain sequence information. Half-life outliers of *S. pombe* (half-life less than 1 or larger than 250 mins) were removed. For both half-life datasets, only mRNAs with mapped 5’UTR and 3’UTR were considered. mRNAs with 5’UTR length shorter than 6nt were further filtered out. Codon-wise species-specific tRNA adaptation index (sTAI) of yeasts were obtained from Sabi and Tuller (Sabi & Tuller, 2014). Gene-wise sTAIs were calculated as the geometric mean of sTAIs of all its codons (stop codon excluded).

### Analysis of knockout strains

The effect level of an individual sequence feature was compared against the wild-type with Wilcoxon rank-sum test followed by multiple hypothesis testing p-value correction (FDR < 0.1). For details see Supplementary methods.

### Motif discovery

Motif discovery was conducted for the 5’UTR, the CDS and the 3’UTR regions. A linear mixed effect model was used to assess the effect of each individual k-mer while controlling the effects of the others and for the region length as a covariate as described previously (Eser *et al*, 2016). For CDS we also used codons as further covariates. In contrast to Eser and colleagues, we tested the effects of all possible k-mers with length from 3 to 8. The linear mixed model for motif discovery was fitted with GEMMA software (Zhou *et al*, 2013). P-values were corrected for multiple testing using Benjamini-Hochberg’s FDR. Motifs were subsequently manually assembled based on overlapping significant (FDR < 0.1) k-mers.

### Folding energy calculation

RNA sequence folding energy was calculated with RNAfold from ViennaRNA version 2.1.9 (Lorenz *et al*, 2011), with default parameters.

### *S. cerevisiae* conservation analysis

The phastCons (Siepel *et al*, 2005) conservation track for *S. cerevisiae* was downloaded from the UCSC Genome browser (http://hgdownload.cse.ucsc.edu/goldenPath/sacCer3/phastCons7way/). Motif single-nucleotide level conservation scores were computed as the mean conservation score of each nucleotide (including 2 extended nucleotide at each side of the motif) across all motif instances genome-wide (removing NA values).

### Linear model for genome-wide half-life prediction

Multivariate linear regression models were used to predict genome-wide mRNA half-life on the logarithmic scale from sequence features. Only mRNAs that contain all features were used to fit the models, resulting with 3,862 mRNAs for *S. cerevisiae* and 3,130 mRNAs for *S. pombe*. Out-of-fold predictions were applied with 10-fold cross validation for any prediction task in this study. For each fold, a linear model was first fitted to the training data with all sequence features as covariates, then a stepwise model selection procedure was applied to select the best model with Bayesian Information Criterion as criteria (*step* function in R, with k = log(n)). L1 or L2 regularization were not necessary, as they did not improve the out-of-fold prediction accuracy (tested with glmnet R package (Friedman *et al*, 2010)). Motif discovery was performed again at each fold. The same set of motifs were identified within each training set only. For details see Supplementary methods.

### Analysis of sequence feature contribution

Linear models were first fitted on the complete data with all sequence features as covariates, non-significant sequence features were then removed from the final models, ending up with 70 features for *S. cerevisiae* model and 75 features for *S. pombe* (each single coding codon was fitted as a single covariate). The contribution of each sequence feature was analyzed individually as a univariate regression and also jointly in a multivariate regression model. The contribution of each feature *individually* was calculated as the variance explained by a univariate model. Features were then added in a descending order of their individual explained variance to a joint model, *cumulative* variance explained were then calculated. The *drop* quantify the drop of variance explained as leaving out one feature separately from the full model. All contributions statistics were quantified by taking the average of 100 times of 10-fold cross-validation.

### Single-nucleotide variant effect predictions

The same model that used in sequence feature contribution analysis was used for single-nucleotide variant effect prediction. For **motifs,** effects of single-nucleotide variants were predicted with linear model modified from (Eser *et al*, 2016). When assessing the effect of a given motif variation, instead of estimating the marginal effect size, we controlled for the effect of all other sequence features using a linear model with the other features as covariates. For details see Supplementary methods. For **other sequence features,** effects of single-nucleotide variants were predicted by introducing a single nucleotide perturbation into the full prediction model for each gene, and summarizing the effect with the median half-life change across all genes. For details see Supplementary methods.

### Construction of *SFG1* and *NYV1* mutant strains

100 bp primers (IDT) containing the respective 3’UTR mutations were used to amplify the kanMX cassette from plasmid pFA6a-KanMX6 (Euroscarf). PCR products were used for transformation of strain BY4741 (MATa his3Δ1 leu2Δ0 met15Δ0 ura3Δ0, Euroscarf) by homologous recombination, and transformants were selected on G418 plates. Correct clones were confirmed by sequencing. Sequences of the constructs are given in Supplementary Table 7.

### Quantitative PCR

Cells were grown to OD_600_ 0.8 in YPD from overnight cultures inoculated from single colonies. Cells were centrifuged at 4,000 rpm for 1 min at 30°C and pellets were flash-frozen in liquid nitrogen. RNA was phenol/chloroform purified. cDNA synthesis was performed with 1.5 μg RNA using the Maxima Reverse Transcriptase (Thermo Fisher). qPCR was performed on a qTower 2.2 (Analytik Jena) using a 2 min denaturing step at 95°C, followed by 39 cycles of 5 s at 95°C, 10 s at 64°C, and 15 s at 72°C with a final step at 72°C for 5 min. qPCR was performed using the SensiFAST SYBR No-ROX Kit (Bioline). Primer efficiencies were determined by performing standard curves for all primer combinations. All primer pairs had efficiencies of 95% or higher. Sequence information of primer pairs and efficiencies are provided in Supplementary Table 7. Ct data from three biological and three technical replicates were used for analysis.

Details of analyzing qPCR data are described in supplementary methods.

### Code availability

Analysis scripts are available at: https://i12g-gagneurweb.in.tum.de/gitlab/Cheng/mRNA_half_life_public.

## FUNDING

JC and ŽA are supported by a DFG fellowship through QBM. JG was supported by the Bundesministerium für Bildung und Forschung, Juniorverbund in der Systemmedizin “mitOmics” (grant FKZ 01ZX1405A).

## ACKNOWLEDGEMENTS

We are thankful to Patrick Cramer for supporting the motif validation experiment. We are thankful to Fabien Bonneau (Max Planck Institute of Biochemistry) for helpful discussions on motifs and RNA degradation pathways, as well as useful feedback on the manuscript. We thank Björn Schwalb for communication on analyzing the knockout data. We thank Vicente Yépez for useful feedback on the manuscript and Patrick Cramer for institutional support.

## 2 Supplementary Figures

**>Figure S1:**
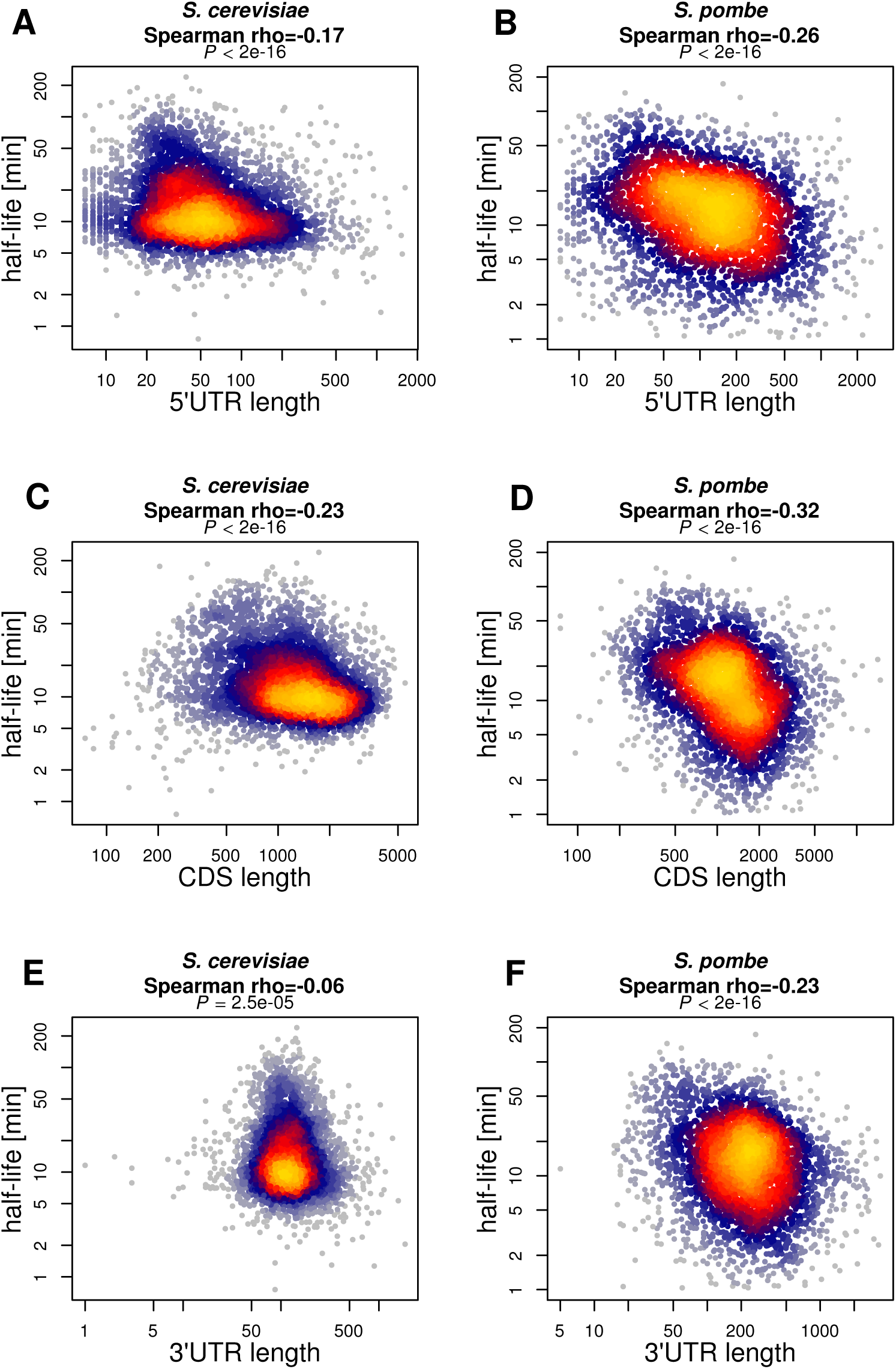
Length of 5*’* UTR, CDS and 3*’* UTR correlate with mRNA half-life. (**A-B**) 5*’*UTR length (x-axis) versus half-life (y-axis) for *S. cerevisiae* (A) and *S. pombe* (B). (**C-D**) CDS length (x-axis) versus half-life (y-axis) for *S. cerevisiae* (C) and *S. pombe* (D). (**E-F**) 3*’*UTR length (x-axis) versus half-life (y-axis) for *S. cerevisiae* (E) and *S. pombe* (F).

**Figure S2:**
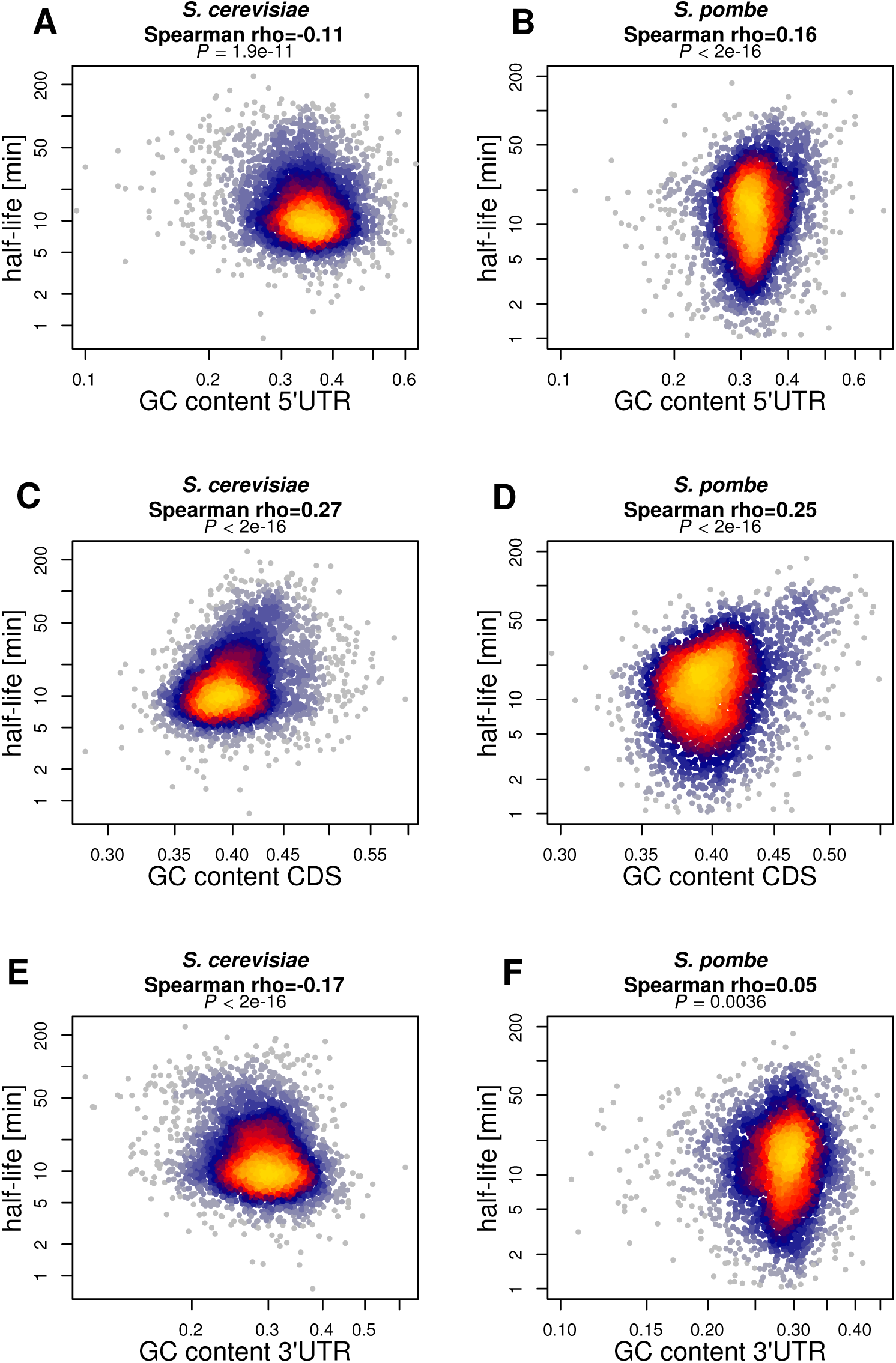
GC content of 5*’*UTR, CDS and 3*’* UTR correlate with mRNA half-life. (**A-B**) 5*’*UTR GC content (x-axis) versus half-life (y-axis) for *S. cerevisiae* (A) and *S. pombe*. (**C-D**) CDS GC content (x-axis) versus half-life (y-axis) for *S. cerevisiae* (C) and *S*.*pombe* (D). (**E-F**) 3*’*UTR GC content (x-axis) versus half-life (y-axis) for *S. cerevisiae* (E) and *S. pombe* (F).

**Figure S3:**
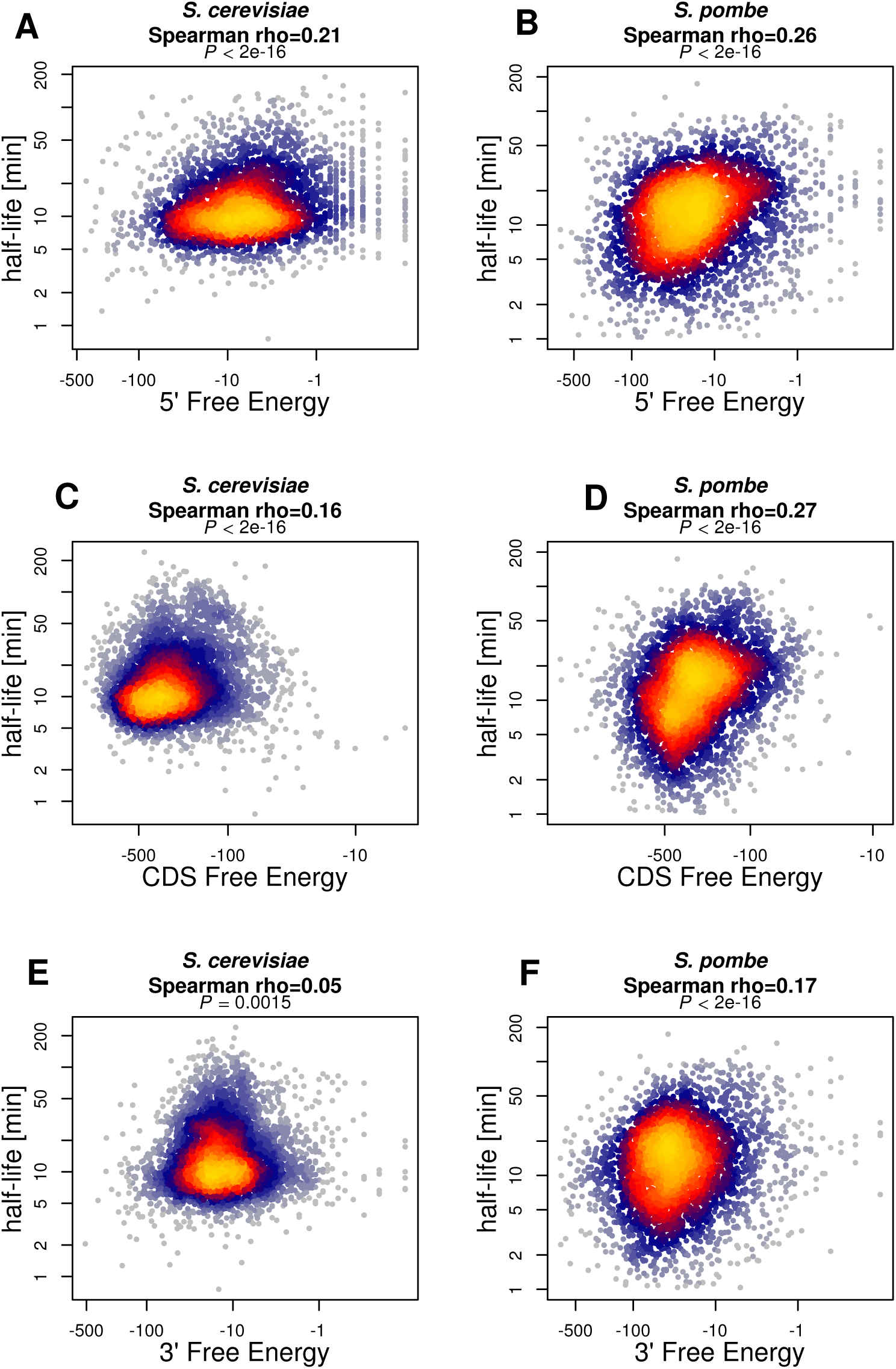
Folding energy of 5*’*UTR, CDS and 3*’*UTR correlate with mRNA half-life. (**A-B**) 5’ free energy (x-axis) lf-versus half-life (y-axis) for *S. cerevisiae* (A) and *S. pombe* (B). (**C-D**) CDS free energy (x-axis) versus half-life (y-axis) for *S.cerevisiae* (C) and *S. pombe* (D). (**E-F**) 3’ free energy (x-axis) versus half-life (y-axis) for *S. cerevisiae* (E) and *S. pombe* (F).

**Figure S4:**
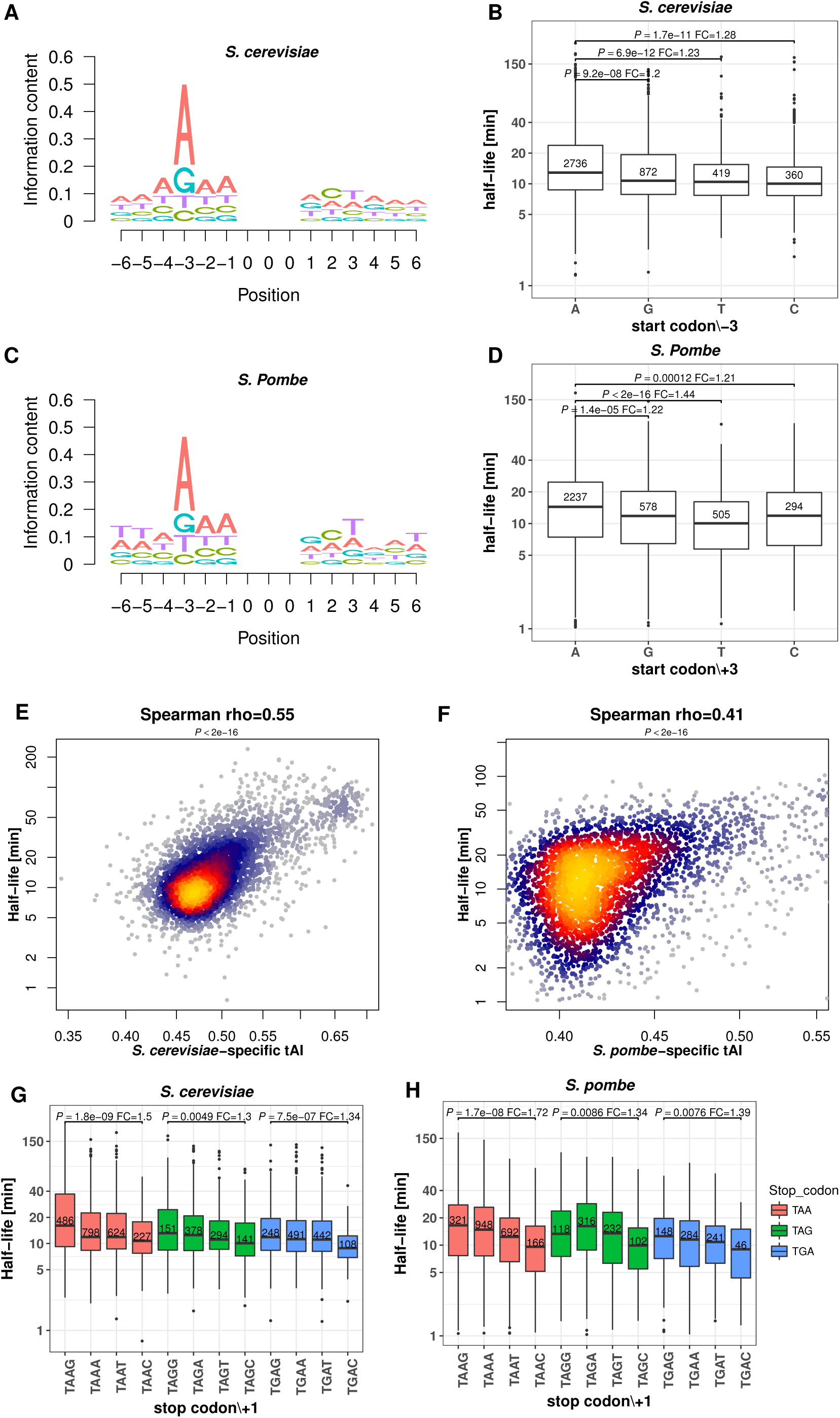
Translation initiation, elongation and termination features associate with mRNA half-life. (**A**) Start codon context (Kozak sequence) generated from 4388 *S. cerevisiae* genes. (**B**) Distribution of half-life for mRNAs grouped by the third nucleotide before the start codon for *S. cerevisiae*. Group sizes (numbers in boxes) show that nucleotide frequency at this position positively associates with half-life. (**C**) Same as (A) for 3713 *S. pombe* genes. (**D**) Same as (B) for *S. pombe*. (**E-F**) mRNA half-life (y-axis) versus species-specific tRNA adaptation index (sTAI) (x-axis) for *S. cerevisiae* (E) and *S. pombe* (F). (**G-H**) Distribution of half-life for mRNAs grouped by the stop codon and the following nucleotide for *S. cerevisiae* (G) and *S. pombe* (H). Colors represent three different stop codons (TAA, TAG and TGA), within each stop codon group, boxes are shown in G, A, T, C order of their following base. Only the P-values for the most drastic pairwise comparisons (A versus C within each stop codon group) are shown. All p-values in boxplot were calculated with Wilcoxon rank-sum test. Boxplots computed as in Figure 3.

**Figure S5:**
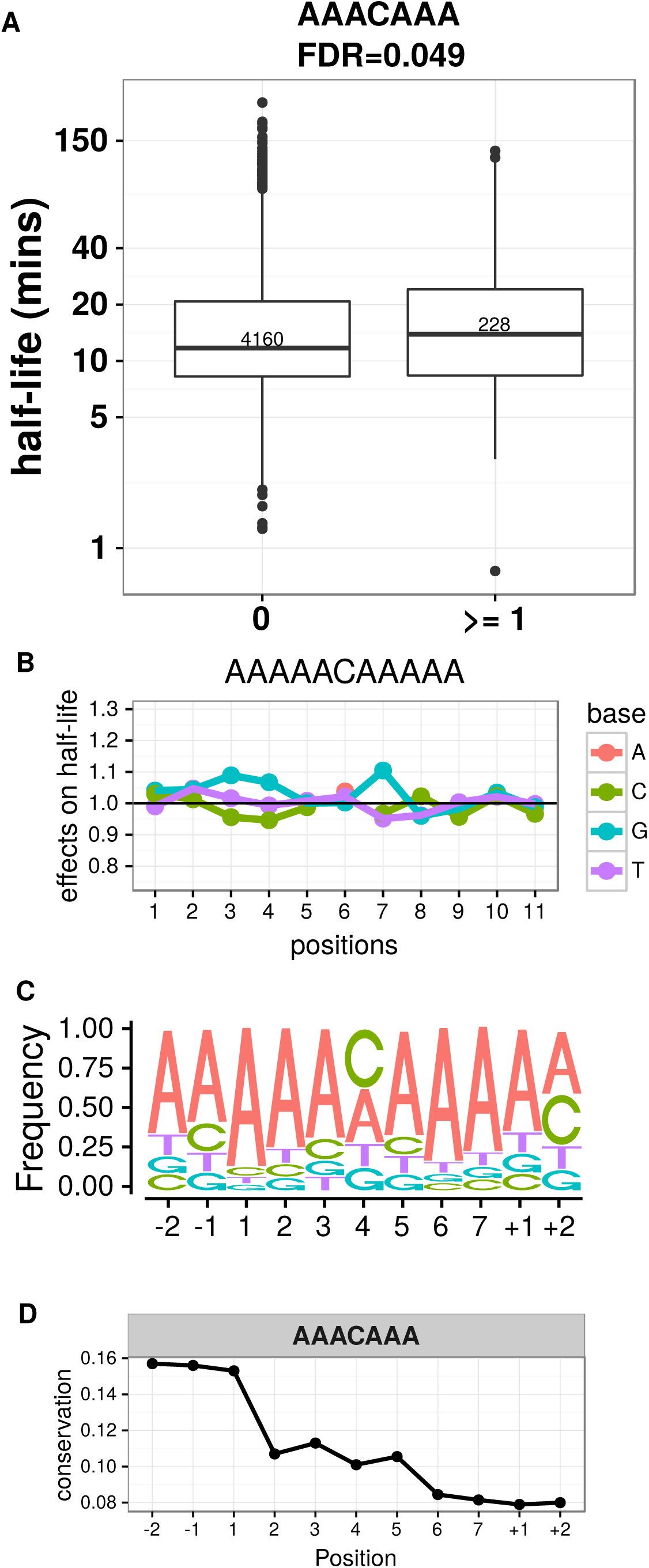
*S. cerevisiae* 5’ UTR mRNA half-life associated motif. (**A**) Distribution of half-lives for mRNAs grouped by the number of occurrence(s) of the motif AAACAAA in their 5’UTR sequence. Numbers in the boxes represent the number of members in each box. FDR were reported from the linear mixed effect model (Materials and Methods). (**B**) Prediction of the relative effect on half-life (y-axis) for single-nucleotide substitution in the motif with respect to the consensus motif (y=1, horizontal line). The motifs were extended 2 bases at each flanking site (positions +1, +2, -1, -2). (**C**) Nucleotide frequency within motif instances, when allowing for one mismatch compared to the consensus motif. (**D**) Mean conservation score (phastCons, Materials and Methods) of each base in the consensus motif with 2 flanking nucleotides (y-axis).

**Figure S6:**
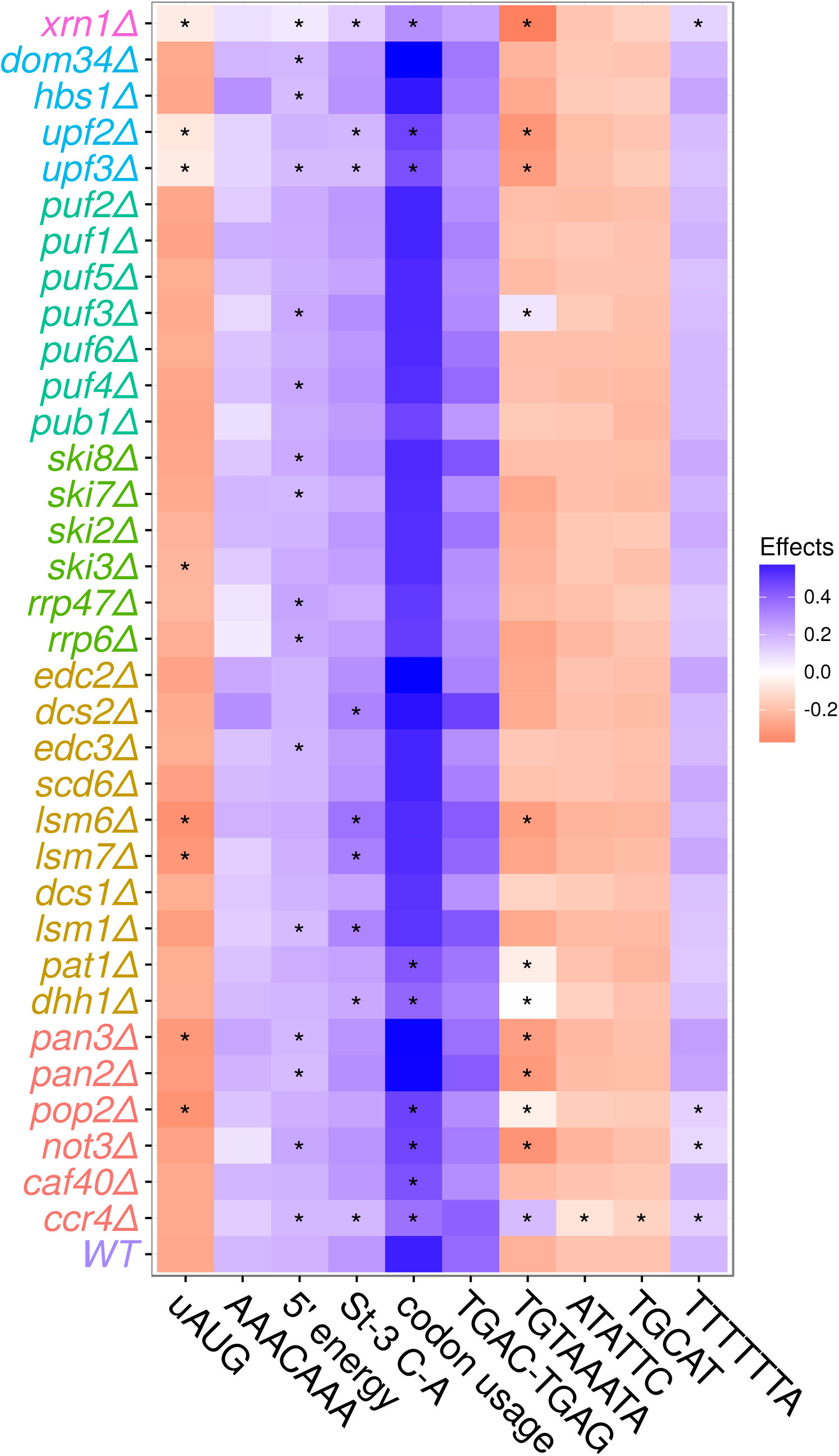
Summary of CREs effect changes across all 34 knockouts comparing with WT. Colour represent the relative effect size (motifs, St-3 C-A, TGAG-TGAC, uAUG), correlation (5’ folding energy) or explained variance (codon usage) upon knockout of different genes (y-axis) (Materials and Methods for detailed description). Wild-type label is shown in the bottom (WT) P-values calculated with Wilcoxon rank-sum test by comparing each mutant to wild-type level, multiple testing p-values corrected with Benjamini & Hochberg (FDR). Stars indicating significance of statistical testing (FDR *<*0.1). 5’ energy: correlation of 5’UTR folding energy with mRNA half-lives; St-3 C-A: relative median half-life difference between genes with cytosine and adenine at start codon -3 position; TGAC-TGAG: relative median half-life difference between genes with stop codon +1 TGAC and TGAG. codon usage: codon usage explained mRNA half-lives variance. uAUG: relative median half-life difference between genes without and with upstream AUG in the 5’UTR (Materials and Methods)

**Figure S7:**
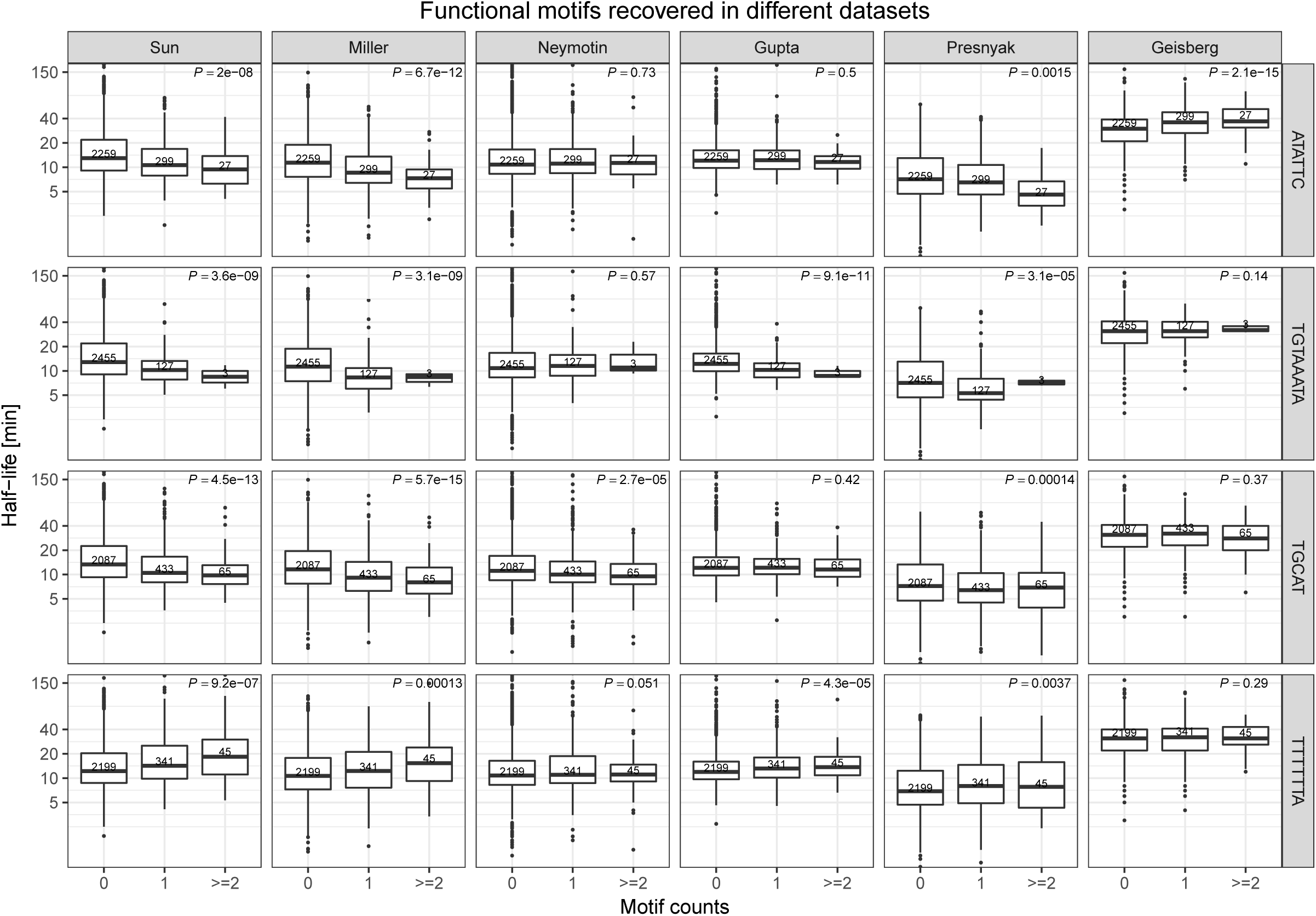
Association of half-life determinant motifs to mRNA half-life in 6 different studies. P-values were calculated with Wilcoxon rank-sum test by comparing half-life of genes without the corresponding motif with genes with the corresponding motif.

**Figure S8:**
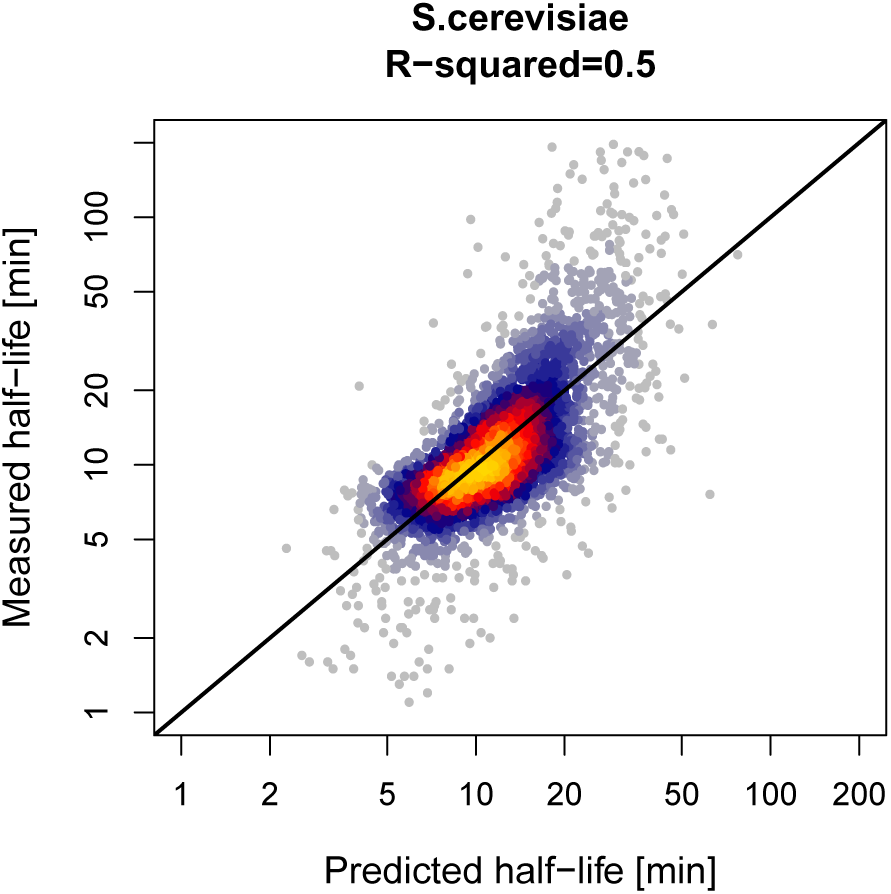
Genome-wide prediction of mRNA half-lives from sequence features with RATE-seq data. mRNA half-lives predicted (x-axis) versus measured (y-axis) with RATE-seq data for 3,539 genes that have complete profiles of all features.

**Figure S9:**
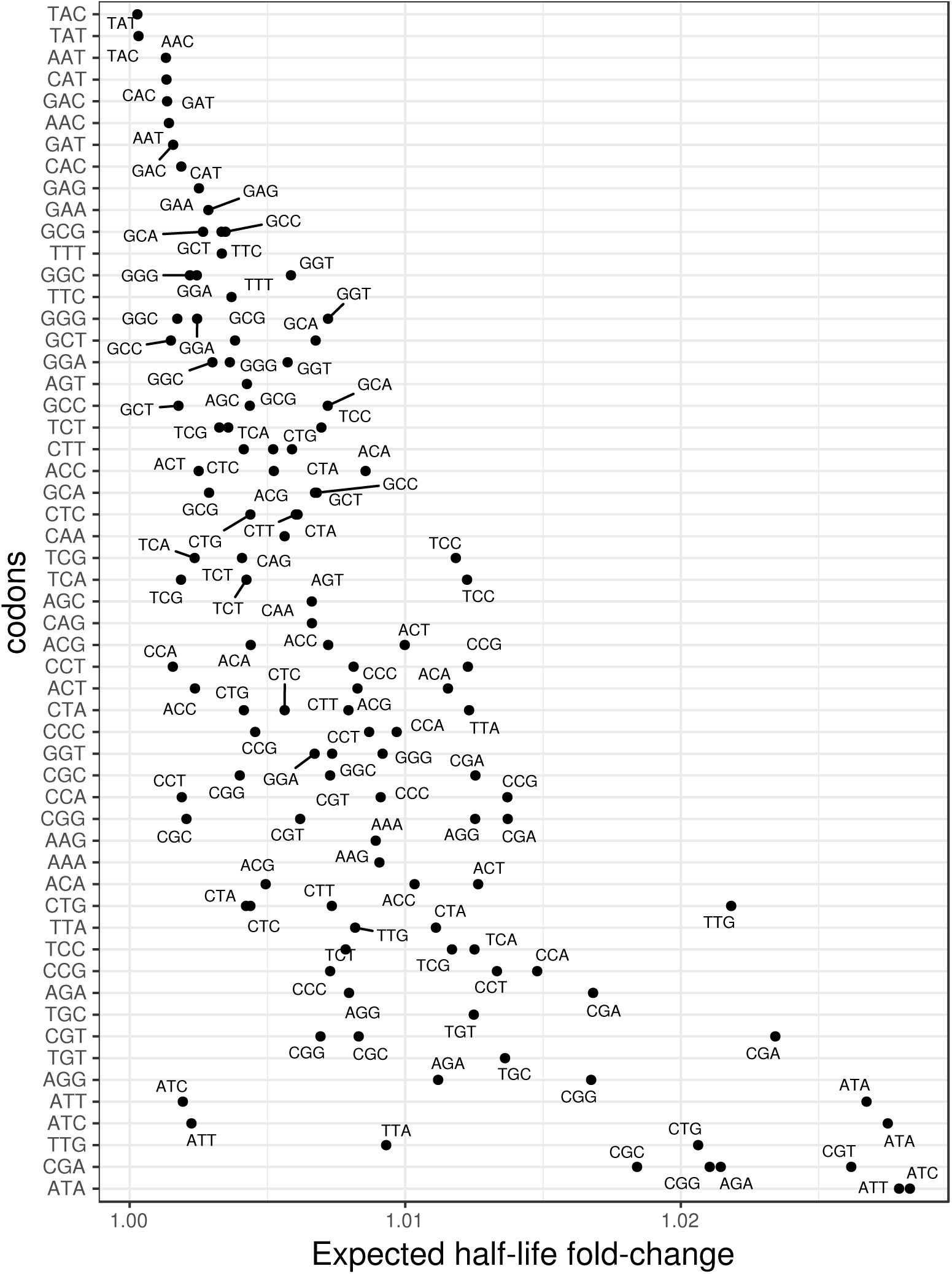
Predicted effects of synonymous codon transitions on half-life. Expected half-life fold-change (x-axis) at each synonymous codon transitions. Each row represent transition from one codon (y-axis) to its synonymous partners. Only synonymous codons that differ by one base were considered.

